# Deciphering the co-evolutionary dynamics of L2 β-lactamases via Deep learning

**DOI:** 10.1101/2024.01.14.575584

**Authors:** Yu Zhu, Jing Gu, Zhuoran Zhao, A W Edith Chan, Maria F. Mojica, Andrea M. Hujer, Robert A. Bonomo, Shozeb Haider

## Abstract

L2 β-lactamases, a serine-based class A β-lactamases expressed by *Stenotrophomonas maltophilia* plays a pivotal role in antimicrobial resistance. However, limited studies have been conducted on these important enzymes. To understand the co-evolutionary dynamics of L2 β-lactamase, innovative computational methodologies, including adaptive sampling molecular dynamics simulations, and deep learning methods (convolutional variational autoencoders and BindSiteS-CNN) explored conformational changes and correlations within the L2 β-lactamase family together with other representative class A enzymes including SME-1 and KPC-2. This work also investigated the potential role of hydrophobic nodes and binding site residues in facilitating the functional mechanisms. The convergence of analytical approaches utilized in this effort yielded comprehensive insights into the dynamic behaviour of the β-lactamases, specifically from an evolutionary standpoint. In addition, this analysis presents a promising approach for understanding how the class A β-lactamases evolve in response to environmental pressure and establishes a theoretical foundation for forthcoming endeavours in drug development aimed at combating antimicrobial resistance.

**Synopsis:** Deep learning is used to reveal the dynamic co-evolutionary patterns of L2 β-lactamases.

- Analysis of hydrophobic nodes and binding site residues provides a detailed understanding of both local and global dynamic evolution, which explain the functional divergences.
- The employment of two distinct deep learning models, the Convolutional Variational Autoencoder (CVAE) and BindSiteS-CNN, facilitates the investigation of conformational shifts, thereby depicting the dynamic evolution of L2 β-lactamases.
- The effectiveness of CVAE and BindSiteS-CNN in dynamic classification is corroborated with selected features.

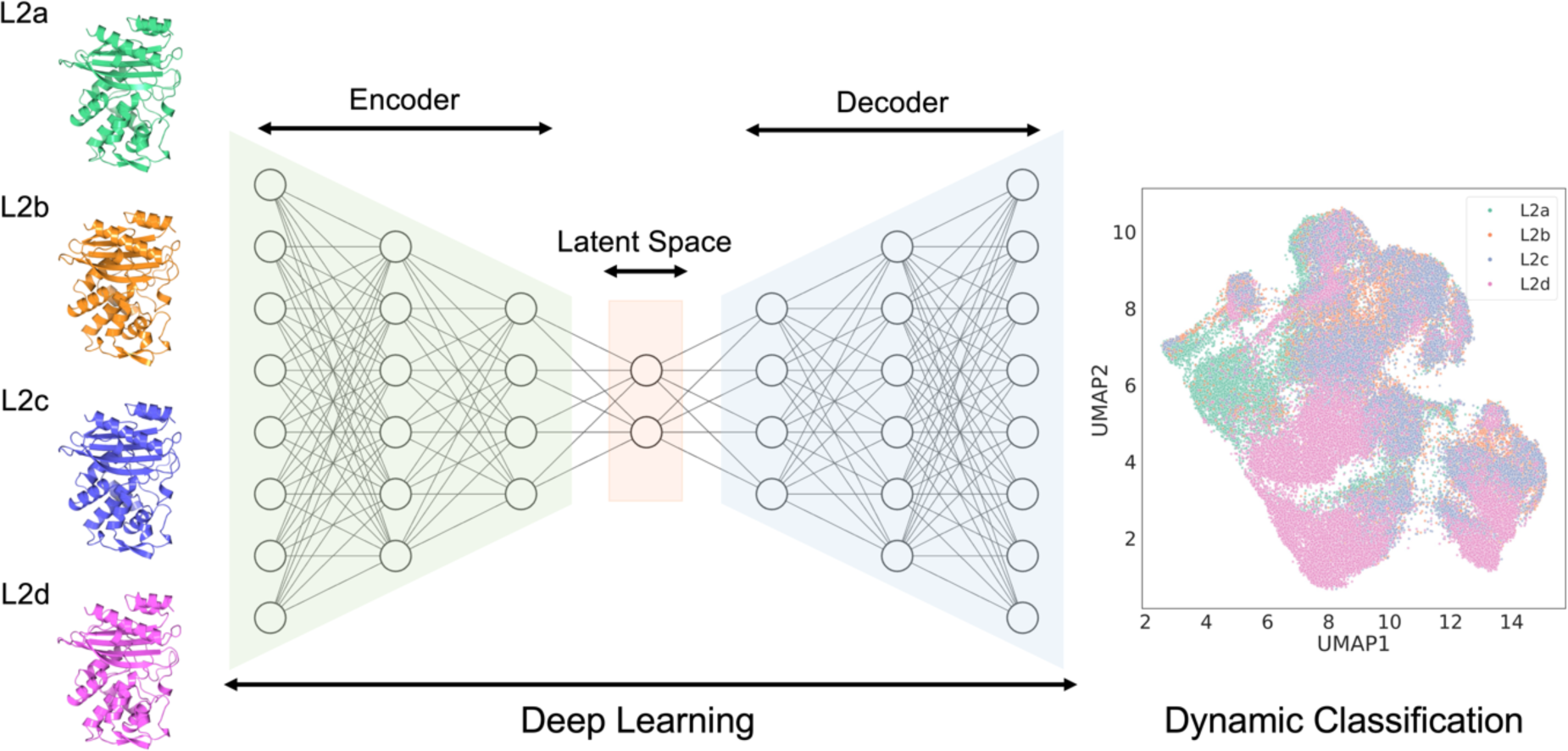

## Introduction

Antimicrobial resistance (AMR) has emerged as a significant public health crisis in the 21st century, threatening a return to the ’pre-antibiotic’ era (Reardon, 2014). This phenomenon, a natural evolutionary response of microbes to antimicrobial agents, risks nullifying decades of medical advancements in treating infectious diseases. Driven by factors such as misuse and overuse of antimicrobials, inadequate infection prevention in healthcare, globalization, and environmental pollution, AMR is growing at an alarming rate. Its clinical implications are dire, with an estimated 1.27 million deaths in 2019 attributed to antibiotic-resistant infections (Perez *et al*, 2014; Murray *et al*, 2022). The potential annual death toll could escalate to 10 million by 2050, alongside a staggering economic loss of 100 trillion USD (Kraker *et al*, 2016).

Central to the battle against bacterial infections are β-lactam antibiotics, characterized by their unique β-lactam ring structure. Their chemical structure is marked by a β-lactam ring, a four-membered cyclic amide. These antibiotics, including penicillins, cephalosporins, carbapenems, and monobactams, revolutionized healthcare since their discovery in the mid-20th century (Bush & Bradford, 2020). The β-lactams function by disrupting bacterial cell wall synthesis, a process vital to bacteria but absent in human cells. They achieve this by targeting penicillin-binding proteins (PBPs), thus leading to bacterial cell death (Bush & Bradford, 2020). However, the extensive and sometimes indiscriminate use of β- lactam antibiotics has accelerated the emergence of bacterial resistance mechanisms. The predominant mechanism involves bacterial β-lactamases, enzymes that deactivate the drug by cleaving the β-lactam ring. Other resistance mechanisms include alterations in PBPs, altered transport into the cell, reduced bacterial membrane permeability to the antibiotic, and enhanced drug expulsion from bacterial cells (Bush & Bradford, 2020).

β-Lactamases are classified into four classes (A, B, C, and D) based on the Ambler molecular classification (Ambler, 1980), and three primary groups based on the Bush-Jacoby-Medeiros functional classification (Bush *et al*, 1995). Each class and group exhibit varying levels of activity against different β-lactam antibiotics, complicating the fight against β-lactamase-mediated resistance (Bush, 2023; Bush & Jacoby, 2010). The persistence and prevalence of β-lactamase genes across various bacterial species underscore the evolutionary advantage conferred by these resistance elements. Through horizontal gene transfer, β-lactamase genes have disseminated across bacterial communities, accelerating the spread of resistance (Fisher *et al*, 2005). Particularly concerning is the emergence of extended-spectrum β- lactamases (ESBLs) and carbapenemases, which hydrolyse a broader spectrum of β-lactam antibiotics (Livermore, 2008). ESBLs represent an evolution of standard β-lactamases, enzymes that bacteria have utilized for millions of years to resist β-lactam antibiotics. However, ESBLs have a more extensive range of action, being capable of hydrolysing not just penicillins but also cephalosporins and monobactams (Bush, 2023). This extended spectrum of activity emerged in the 1980s, marking a crucial evolutionary adaptation in response to the extensive use of β-lactam antibiotics in human and veterinary medicine.

The major clinical concern with β-lactamase production is the concomitant development of multidrug resistance, further exacerbated by co-resistance mechanisms. The broadening substrate profiles of newer β-lactamases threaten the efficacy of last-resort antibiotics, such as carbapenems and ceftazidime-avibactam (Drawz Sarah & Bonomo Robert, 2010; Tooke *et al*, 2019a). β-lactamase inhibitors, like clavulanic acid, sulbactam, and tazobactam, represent a successful approach to circumvent β-lactamase-mediated resistance (Bush & Bradford, 2020; Tooke *et al*, 2019a). These molecules are structurally similar to β-lactam antibiotics and act as ’suicide inhibitors’ by binding to the enzyme active site, thereby preventing antibiotic hydrolysis. However, the emergence of inhibitor-resistant β-lactamases necessitates the development of next-generation inhibitors (Tooke *et al*, 2019a). Therefore, detailed understanding of β-lactamase function, classification, and evolution is critical for effective antibiotic design and future drug development.

β-lactamases represent a fascinating yet daunting facet of bacterial adaptability. Their rapid evolution and broadening resistance profiles underline the urgency of more extensive research (Brook, 2004).

*Stenotrophomonas maltophilia*, a Gram-negative bacterium, is increasingly recognized as a significant opportunistic pathogen, especially prevalent in hospital environments where it poses a serious risk to immunocompromised patients (Akova *et al*, 1991; Avison *et al*, 2001; Hu *et al*, 2008). The clinical relevance of this pathogen is heightened due to its production of two chromosomally encoded β- lactamases: L1, a metallo-β-lactamase, and L2, a class A cephalosporinase (Avison *et al*, 2001). These enzymes play a pivotal role in conferring resistance to a broad spectrum of β-lactam antibiotics, rendering infections by *S. maltophilia* particularly difficult to treat. The L1 β-lactamase, characterized as a metallo-β-lactamase, relies on zinc ions in its active site for the hydrolysis of β-lactam antibiotics, effectively neutralizing their antibacterial activity (Okazaki & Avison, 2008). Conversely, L2, identified primarily as a class A cephalosporinase, is susceptible to inhibition by clavulanate, a known irreversible inhibitor of class A β-lactamases (Okazaki & Avison, 2008). Both L1 and L2 are inducible enzymes, whose expression is modulated by specific regulatory genes, some of which are shared with those controlling the production of AmpC-type β-lactamases (Hu *et al*, 2008). L2 β-lactamases, have not been extensively studied in spite of them being integral to comprehending antibiotic resistance mechanisms.

Interestingly, despite the shared regulatory genes, the response to induction is distinct for L1 and L2 β- lactamases. This disparity in inducibility adds complexity to the resistance mechanisms employed by *S. maltophilia* (Chang *et al*, 2015). As a result, infections caused by this bacterium are challenging to treat with conventional β-lactam antibiotics, leading to increased mortality rates among affected patients (Brooke, 2012). The prevalence of *S. maltophilia* infections in the hospital setting has been on the rise, and this bacterium is increasingly recognized as a formidable multidrug-resistant pathogen (Mojica *et al*, 2023). Traditionally, *S. maltophilia* has demonstrated resistance to a wide range of β-lactam antibiotics, including penicillins, cephalosporins, and carbapenems. This resistance is primarily attributed to the expression of L1 and L2 β-lactamases.

Recent investigations into β-lactam–β-lactamase inhibitor combinations have uncovered promising options. One such combination, aztreonam added to ceftazidime-avibactam, shows potential due to aztreonam’s resistance to L1 hydrolysis and avibactam’s effective inhibition of L2 (Calvopiña *et al*, 2017; Nicodemo & Paez, 2007; Mojica *et al*, 2017). This discovery offers hope in the fight against *S. maltophilia* infections. A pioneer genetic study analysed 130 U.S. clinical isolates of *S. maltophilia*, uncovering significant diversity with 90 different sequence types (Mojica *et al*, 2019). The authors identified 34 novel variants of L1 β-lactamase and 43 novel variants of L2 β-lactamase. Further research has shown that avibactam and a bicyclic boronate can inhibit serine β-lactamase L2, though not metallo β-lactamase L1 (Tooke *et al*, 2019b). X-ray crystallography provided insights into the binding mechanics of these inhibitors, revealing covalent bonding with L2’s nucleophilic serine. The resolution of the apo structure of L2b (PDB ID: 5NE2, Figure 1A) and in co-complex with avibactam (PDB ID: 5NE3) and the bicyclic boronate 2 (PDB ID: 5NE1) were determined to be 1.19 Å, 1.35 Å and 2.09 Å, respectively (Calvopiña *et al*, 2017). Clear Fo-Fc electron density maps offered compelling evidence that both inhibitors form a covalent bond with the active site nucleophile, serine. Analysis of the binding dynamics of both compounds with L2 revealed no significant morphological changes in its active site when compared with either the apo or the D-glutamate structures. Intriguingly, in the structures of both complexes, the deacylating water molecule is positioned analogously to its placement in the native and D-glutamate-bound structures (Calvopiña *et al*, 2017), thereby suggesting a preserved catalytic mechanism in the active site of L2, despite the inhibitors’ covalent binding (Tooke *et al*, 2019b).

**Figure 1.**
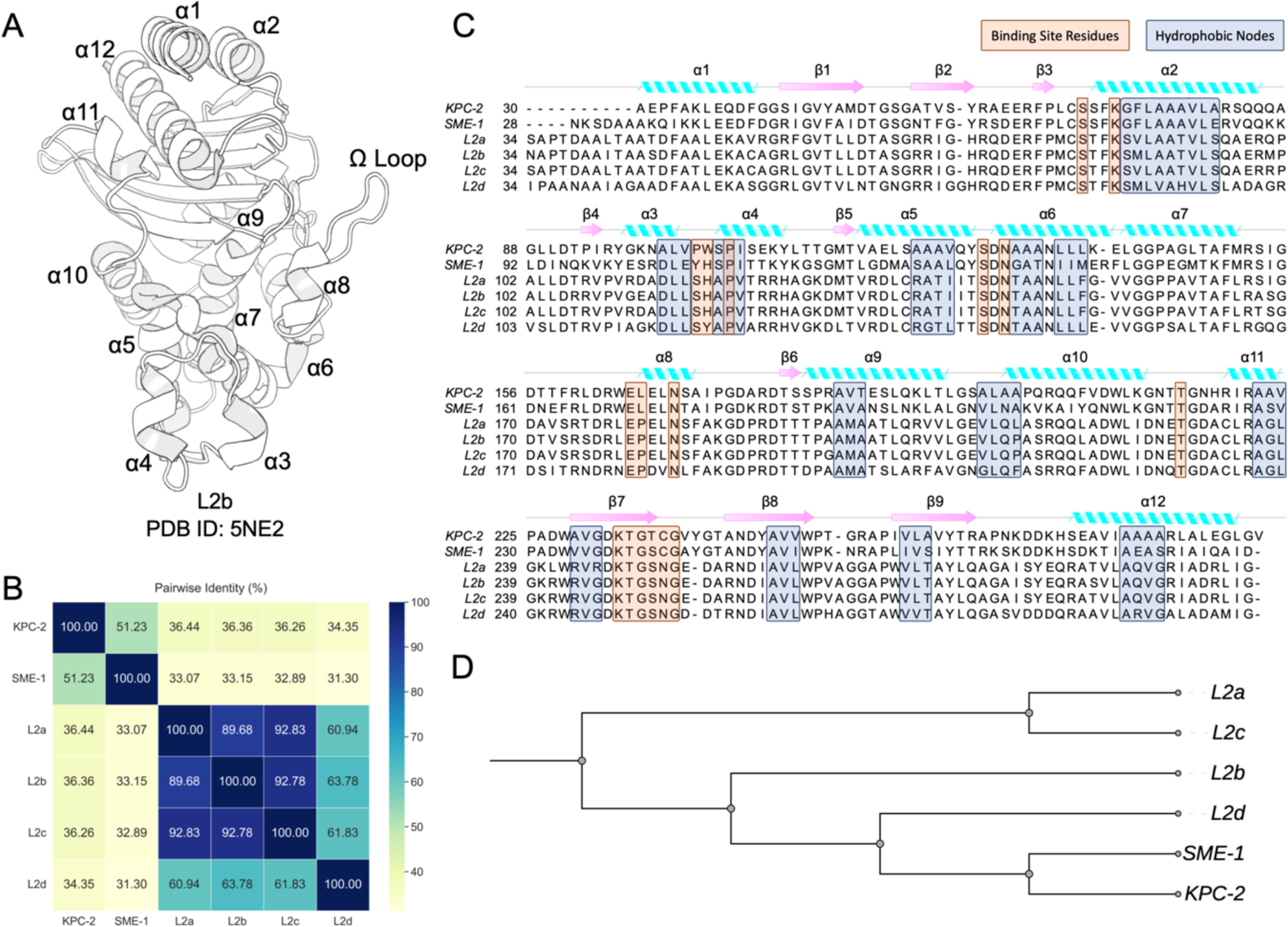
Evolution of L2 β-lactamases. (A) Crystal structure of L2b β-lactamase (PDB ID: 5NE2) (Calvopiña et al., 2017). (B) Pairwise sequence identity of the six class A β-lactamase systems. (C) Multiple sequence alignments between KPC-2, SME-1, L2a, L2b, L2c and L2d with secondary structure element annotations. The blue boxes indicate the hydrophobic nodes while orange boxes represent binding site residues. (D) Phylogenetic tree of six class A β-lactamase systems based on the multiple sequence alignment.

The conducted genetic investigation certainly enriched the knowledge on the pervasiveness and variety of L2 β-lactamase (Walsh *et al*, 1997). However, it compels us to provide indispensable structural and functional information about the repercussions of the identified amino acid changes. Such information would offer profound insight into the protein’s activity and behaviour. Additionally, the absence of a deep understanding of the dynamic consequences these allelic variations bring about presents a noteworthy gap in the study. These dynamics, which illustrate how proteins behave and change over time, are vitally important when attempting to predict the molecular-level impacts of these variants. The study included novel L2 genes L2b, L2c, and L2d from isolates K279a, J675a, and N531 while the L2a was sequenced from strain IID 1275 L2 (Avison *et al*, 2001; Mojica *et al*, 2019). Their sequences differ from the published sequence of strain IID 1275 L2 (L2a) by 9%, 4%, and 25% respectively, highlighting considerable strain divergence. Interestingly, this extensive sequence variation was not mirrored in the differences between the 16S rRNA genes from the same isolates, where genetic drift was less than 1% (Avison *et al*, 2001). The observed variation among the L2 β-lactamase genes might suggest accelerated evolution, implicating horizontal gene transfer in the process.

Apart from the sequence and structure differences, the co-evolutionary dynamics offer new insights into how homologous β-lactamases respond to residue substitutions, suggesting that incorporating this information into the development of β-lactamase inhibitors could lead to the creation of more effective drugs with broader applications. This research focuses on the dynamic co-evolution of L2 β-lactamases and other representative class A β-lactamases (SME-1 and KPC-2), leveraging molecular simulations and deep learning methods. Specifically, this work aims to differentiate and categorize conformations through the synergistic use of convolutional neural networks and autoencoders, integrating co-evolution information into the representation of protein dynamic properties.

To examine the conformational landscape of the six β-lactamase systems, adaptive bandit simulations were run (Perez *et al*, 2020). The dynamic similarity of these systems was assessed using unsupervised low-dimensional embeddings, derived from a convolutional variational autoencoder (CVAE) (Bhowmik *et al*, 2018) and BingSiteS-CNN (Scott *et al*, 2022). Insights from these deep learning techniques revealed critical conformational changes and similarities, impacting the dynamic architecture of the enzyme’s active site and potentially its catalytic activity evolution. Additionally, the research explores the conservation and specificity of the sequence, structure, and dynamic properties of β-lactamase to uncover correlations with enzymatic functions, thereby constructing a comprehensive sequence-structure-dynamics-function hypothesis landscape.

Our findings offer a detailed understanding of the behaviour of L2 β-lactamases and two other well-studied representative class A β-lactamases (SME-1 and KPC-2) from an evolutionary perspective. This knowledge is vital for rational drug discovery, assisting in the design of drugs that effectively modulate protein behaviour. By addressing the knowledge gap in describing the evolution pathway using dynamics of variations, this work not only enhances understanding of L2 β-lactamase variants but also aims to drive innovation in drug discovery.

## Results

### Sequence and structure evolution

The six studied enzymes belong to class A β-lactamases according to the Ambler classification, which is based on homology. However, there is variation in their pairwise identities ranging from 31.30% to 92.83% (Figure 1B). A comprehensive L2 β-lactamase alignment of the sequence length found a little disparity, whereby L2a, L2b, and L2c exhibited a length of 270 residues; however L2d exhibited an extra residue, namely G271, located inside the β2 sheet. The observed deviation in L2d might potentially have consequences for the structure and function of proteins. Nevertheless, the exact implications of this variation remain uncertain and require further inquiry for achieving a more complete understanding.

A total of 13 hydrophobic nodes, consisting of 48 residues, were identified during the alignment (Figure 1C). The spatial analysis revealed that a majority (10 out of 13) were located inside α-helices, while the other three were situated in the β sheets. Consistent with what has been observed in other class A β- lactamases, the α- and β-hydrophobic network were also present in L2 family members (Galdadas *et al*, 2018; Olehnovics *et al*, 2021). Significantly, these nodes exhibited inter-nodal interactions, namely packing, with one another, hence enhancing the flexibility and robustness of the tertiary structures. Meanwhile, the binding site residues comprised of 17 residues, one of which, a proline located in the α4 helix, coincided with the hydrophobic nodes. These residues were mostly found in the α domain, with a few at the interface of the α and α-β domains. They constitute the essential core of the active site and play a direct role in enzymatic activity.

The phylogenetic analysis (Figure 1D) highlighted a discernible pattern in the evolutionary connections among the L2 β-lactamases, SME-1 and KPC-2. The constructed phylogenetic tree posits SME-1 and KPC-2 as the evolutionarily closest enzymes to L2d, suggesting a relatively common ancestor amongst these three β-lactamases. Following SME-1 and KPC-2, the L2b variant is identified to bear a closer phylogenetic relationship with L2d rather than L2a and L2c, thereby indicating a particular evolutionary bifurcation. This observed association suggests that L2d may have undergone divergence from a common ancestor concomitantly shared with SME-1 and KPC-2, prior to its divergence from L2a and L2c, with L2b assuming an intermediary stance within the evolutionary trajectory. Nonetheless, the precise selective pressures precipitating these evolutionary patterns necessitate further investigation.

Pairwise structural alignment of the initial structures of six systems resulted in an average Cα atom root-mean-square deviation (RMSD) of 0.65Å (Figure S1). Particularly, within the L2 family, the average RMSD value was only 0.10Å. Figure 1C illustrates the alignment differences of each pair among the six β-lactamases. It should be highlighted that the secondary structure elements, such as α helices and β sheets, displayed strong alignment, indicating conserved structural properties across the sequence evolution. In addition, important regions such as the conformation of the Ω loop, the hinge region, and the loop between α3 and α4 helix were conserved. The preservation of these regions indicates their functional importance or structural stability, and it suggests their crucial roles in the protein’s overall functions and stability across different systems.

### Dynamics evolution identified by CVAE

An unsupervised Convolutional Variational Autoencoder (CVAE)-based deep learning approach was used to investigate the conformational transitions triggered by alterations in interactions (Figure S2). It is particularly adept at handling the complex and dynamic 3D structures of proteins due to its unique architecture. By taking the advantages of both convolutional neural network and variational autoencoder, CVAE has demonstrated significant success in diverse applications such as protein folding analysis (Bhowmik *et al*, 2018), investigating glycosyltransferase (Akere *et al*, 2020) and glucocerebrosidase (Romero *et al*, 2019) activation and dysfunction, understanding the mechanisms of SARS-CoV-2 (Cho *et al*, 2021), studying how hydrophobic nodes affect class A β-lactamases (Olehnovics *et al*, 2021), and specifically looking into the L1 β-lactamase (Zhao *et al*, 2023). This study focused on identifying dynamic changes and comparing the systems to discern a co-evolutionary pattern. The dynamics involving the hydrophobic nodes and binding site residues were characterized by using a symmetric distance matrix of dimensions 64 × 64, which represented the distances between the featured Cα residues. Before training the CVAE model, the trajectories from the four L2 systems were stacked together as the combined dataset. The model selected for decoding was the latent dimension constructed with the lowest loss, which was 21st dimension (Figure S3). Convergence of the model was attained after the 95th epoch (Figure S3), and the model generated at this epoch was utilised for further decoding.

A pair of concomitant experiments were systematically run in an endeavour to reconstruct the embeddings in parallel. The first experiment concentrated on the decoding process, employing solely the L2 β-lactamase information. The comparison between the origin distance matrix and the decoded matrix revealed a significant similarity, providing evidence that the model effectively captured the essential characteristics from both the distance matrix and the trajectories without losing much useful information (Figure S3). Upon subjecting this data to a rigorous compression and subsequent dimensionality reduction, the elucidated output manifested the conformational clustering trend to the L2 variants. Figure 2 illustrates an intricate delineation of the L2 trajectories within the dimension-reduced simulations.

**Figure 2.**
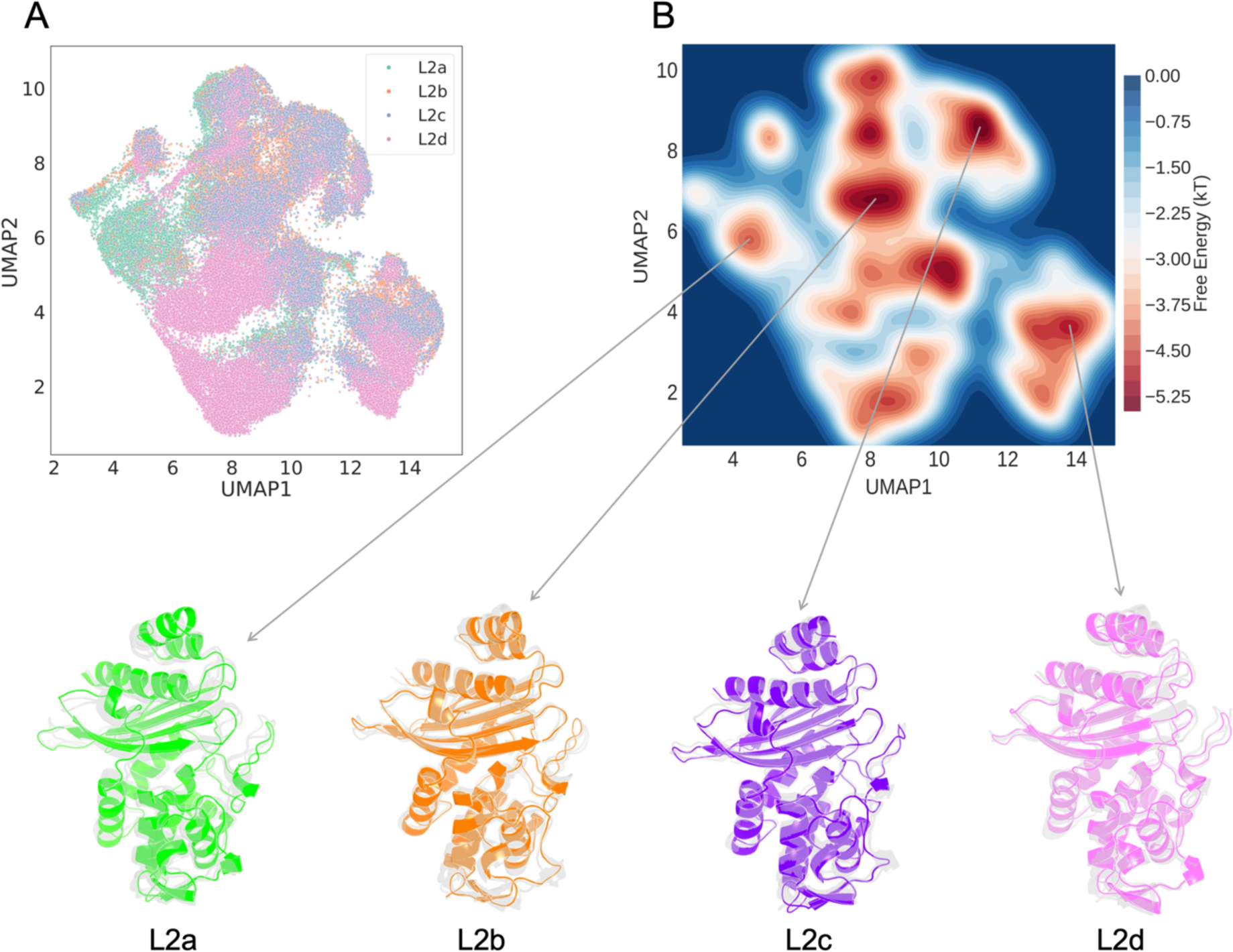
CVAE-based deep clustering of L2 β-lactamase. (A) 2D UMAP projection of the high-dimensional embeddings of L2a (green), L2b (orange), L2c (amethyst) and L2d (violet). (B) The free energy landscape observed within the stacked simulations. Conformations were extracted from the different energy basins for L2a, L2b, L2c and L2d, respectively.

The free energy landscape was generated to represent the distribution of dimensions within the UMAP latent space, providing an initial approximation for cluster identification. Regions of lower energy were indicated by a red hue. The distribution of the system was subsequently projected onto the free energy profile, facilitating the extraction of diverse conformations from the energy basins.

The distinct clustering within the L2 systems, notably that the L2d cluster is markedly discrete compared to the other three counterparts (Figure 2). Furthermore, a portion of the L2a cluster exhibits unique characteristics when compared to the remaining clusters. The application of the CVAE architecture successfully grouped conformations by selected features, confirming its utility in clustering.

To substantiate the usefulness of this clustering, multiple conformations from the free energy basins were extracted for each L2 system. Despite variations in the sequences across different systems, the principal structures remained consistent throughout the evolutionary process. Minor discrepancies in the α- and β-networks were observed; however, the core structures, including key binding site residues, are almost identical. This consistency highlights the adaptive function of these residues in preserving the tertiary structure of the L2 β-lactamase and the effectiveness of hydrophobic nodes and binding site residues in describing the dynamics evolution.

The Ω loop, the α3-α4 loop, and an additional hinge-region loop, exhibited enhanced flexibility relative to the stable core. These dynamic regions play a pivotal role in the functional diversity of β-lactamases, potentially providing insights into the functional variations triggered by sequence differences.

Upon integrating dynamics data from other representative class A β-lactamases like SME-1 and KPC- 2, the second experiment utilized a CVAE model from prior research (Olehnovics *et al*, 2021), decoding the embedding with additional class A β-lactamase trajectories. System distribution analysis, as shown in Figure 3, revealed distinct clustering patterns among the six systems. UMAP plot markers represented unique conformations, fundamental to the model’s encoding and decoding processes. SME-1 and KPC- 2 were distinctly separated from the L2 family, attributable to their carbapenemase activity, in contrast to the ESBL cephalosporinase classification of L2 β-lactamases. Some SME-1 and KPC-2 conformations appeared within the L2d cluster, suggesting a closer similarity to L2d than to other L2 variants. The plot’s proximity metrics indicated this similarity, supporting the hypothesis of L2d β- lactamase’s dynamical closeness to SME-1 and KPC-2. This aligns with sequence phylogenetic findings. Specifically, the SME-1 cluster was near L2 variants than KPC-2, implying a closer dynamic relationship with L2 β-lactamase. Consequently, the evolutionary trajectory from the dynamics perspective for class A β-lactamase appears as KPC-2 ® SME-1 ® L2d ® L2a/b/c.

**Figure 3.**
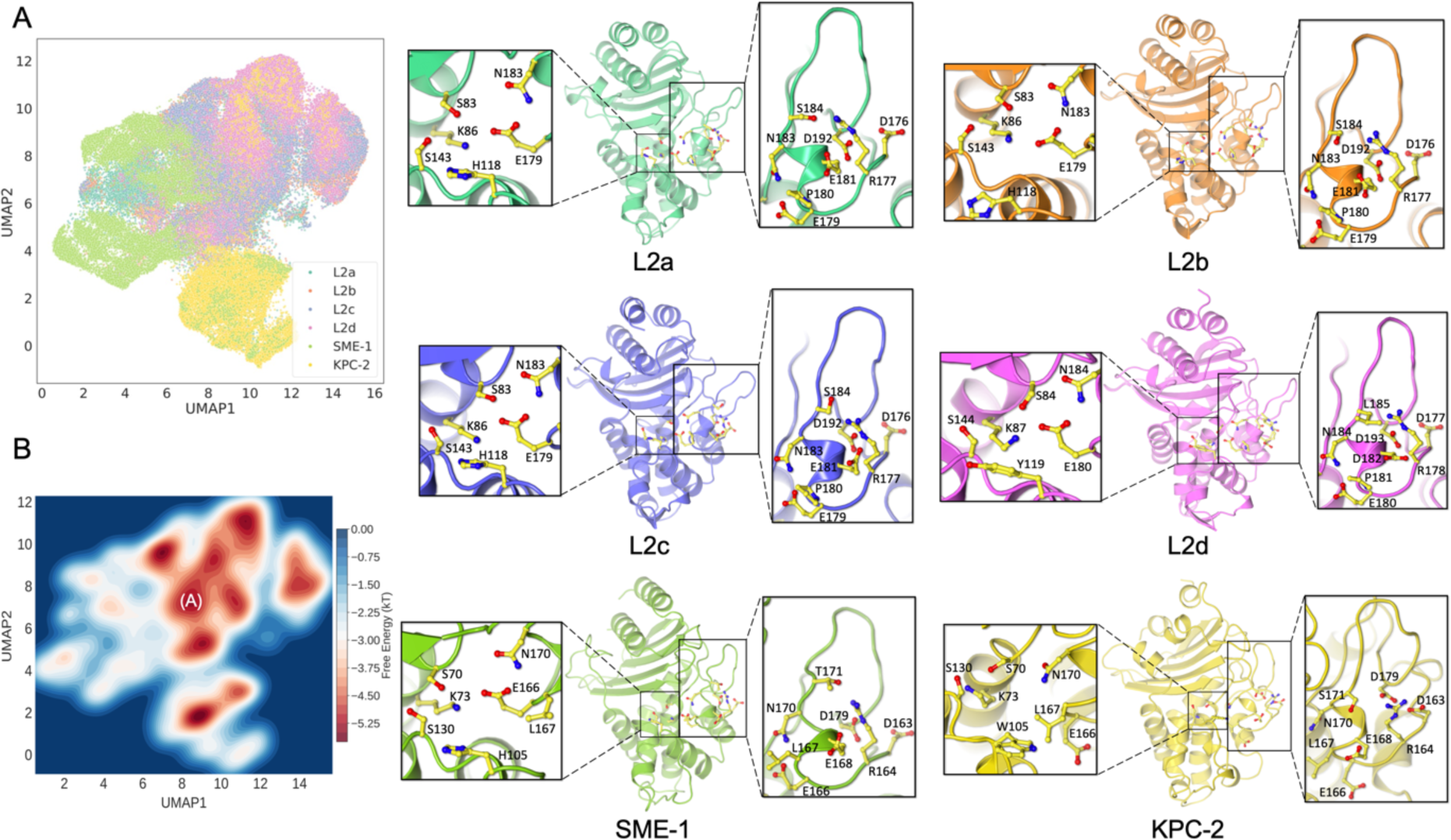
CVAE-based deep clustering of the six class A β-lactamase. (A) 2D UMAP projection of the high-dimensional embeddings of L2a (green), L2b (orange), L2c (amethyst), L2d (violet), SME-1 (pale green) and KPC-2 (yellow). (B) Histogram approximation of the free energy landscape observed within the stacked simulations. Structures from basin A were extracted for local dynamics investigation. Conformations were extracted from the energy basins for L2a, L2b, L2c, L2d, SME-1 and KPC-2 respectively. Local details of the active site and the essential Ω loop are highlighted.

Further examination involved extracting structures from the approximated free energy basins of the six class A systems to seek residue-level evidence. Here, structures for each system were extracted from defined basin A in the free energy landscape. The active site, crucial for catalysis, resides in a sub- domain cleft, defined by the Ω loop, the α3-α4 loop, and the hinge-region loop. Figure 3 illustrates the spatial relationships of these residues across different systems. Stability was a constant feature across all systems, particularly in core structures like helices and sheets, with hydrophobic nodes maintaining their conformations without significant shifts. However, the loops around the active site underwent vital structural transformations, altering the conformation of the active site.

Analysis of the aromatic residue, pivotal in the active site (W105 in KPC-2, H105 in SME-1, Y119 in L2d, H118 in L2a/b/c), revealed a trend across all six β-lactamases. W105 in KPC-2 adopted an outward pose, while H105 in SME-1 shifted inward, pointing towards the active site core. Y119 in L2d, with a six-membered ring, was spatially akin to SME-1 but slightly outwardly oriented. H118 in L2b was further from the active site, and in L2a/c, it was closer, resembling the orientation in SME-1. The orientation of the aromatic residue on the α3-α4 loop in L2d was distinct from that in SME-1, L2a/b/c, and KPC-2. This also reflected the effect of this aromatic residue differs on enzymatic dynamics and functions, which is consistent with previous class A β-lactamase studies including KPC-2 , SME-1, SHV-1, and TEM-1 (Papp-Wallace *et al*, 2010; Olehnovics *et al*, 2021; Ke *et al*, 2007).

Focusing on other essential active site residues and those forming the Ω loop revealed a notable pattern. The Ω loop, crucial for substrate binding and catalysis, can undergo specific substitutions that modify enzyme structure and function (Banerjee *et al*, 1998). In KPC-2, this loop interacted closely with the helices and the β-lactamase core, creating a large but shallow active site due to limited residue interactions. In SME-1, the loop was slightly more distant from the helices, with active site residues moving closer and forming a smaller site. H105 played a role in controlling the site’s openness. In L2d, the loop was further apart compared to SME-1, with active site residues and the α8 helix forming a more defined cavity through enhanced interactions. In L2a/b/c, the loop showed minor fluctuations, with strengthened interactions between active site residues. The consistent trend across the six systems, irrespective of residue variations, indicated a progressive movement of active site residues towards each other, coupled with a stretched Ω loop and a consequentially smaller yet deeper active site, potentially influencing selective ligand binding in various β-lactamases.

The deconstruction of the CVAE latent dimensions in this context substantially augmented the evolution depth, thereby enabling the UMAP representation to precisely explain how these class A β- lactamase systems evolved. The deep learning-based analyses implies that the sequence evolution in the β-lactamase enzyme may result in corresponding changes in its structure and dynamics. Understanding these relationships is crucial for comprehending the enzyme’s activity from a holistic approach that incorporates sequence, structure, dynamics, and function.

### BindSiteS-CNN based binding site comparison

BindSiteS-CNN is a Spherical Convolutional Neural Network model trained to analyse the similarity of protein binding sites based on their local physicochemical properties (Scott *et al*, 2022). It has shown the capacity of large-scale inter- and intra-group analysis of protein families on Protein Kinases. Here BindSiteS-CNN was used to reveal local similarities/differences within the binding sites, between the six studied β-lactamases. Our hypothesis was driven based on using UMAP to visualize the descriptors from the BindSiteS-CNN model and the enzymes with similar structural features in the binding site would cluster together.

The defined cluster of binding site residues is identical for L2a, L2b and L2c. The main local difference between L2a/b/c and L2d comes from the residue between α3 and α4 (Figure 4A). With Y119 instead of H118 at this position, L2d has a more hydrophobic active site. The UMAP of BindSiteS-CNN based embeddings highlights the difference between the local features in the binding site of L2a/b/c and L2d as well as the similarities within that of L2a/b/c (Figure 4B).

**Figure 4.**
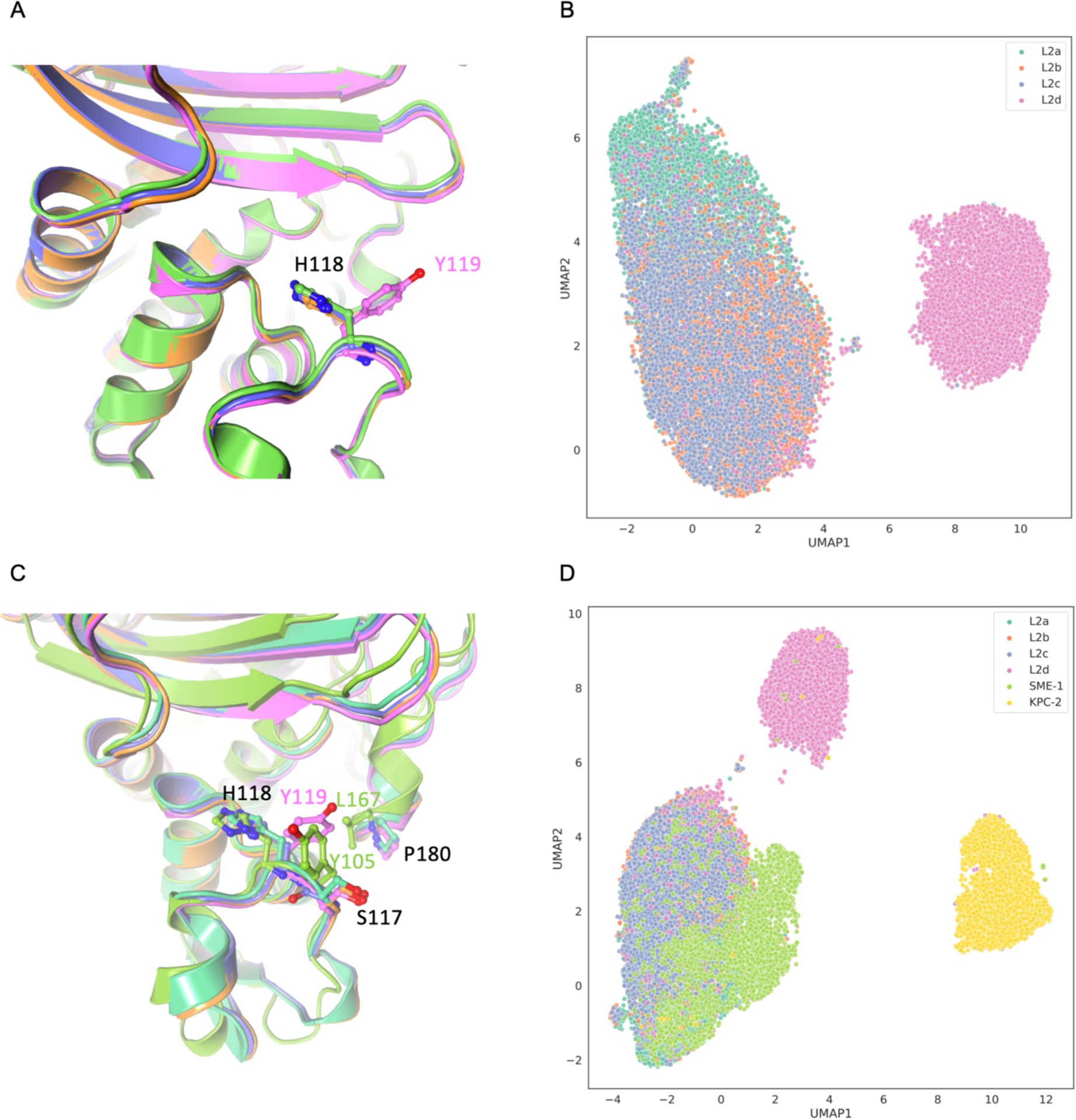
Binding site comparison of L2a (green), L2b (orange), L2c (amethyst), L2d (violet), SME-1 (pale green) and KPC-2 (yellow). (A) The main different residue at the same loci within the defined binding site is H118 of L2a/b/c and Y119 of L2d. (B) BindSiteS-CNN based high-dimensional embeddings represented in 2D with UMAP. (C) Three main different residues within the binding site of L2a/b/c, L2d and SME-1. (D) BindSiteS-CNN based high-dimensional embeddings of all six systems represented in 2D with UMAP.

The UMAP of BindSiteS-CNN based embeddings emphasises the structural differences observed in the local features of the binding site of the six systems with three main clusters (Figure 4D). The main cluster containing L2a/b/c and SME-1 indicates the local similarity between the four systems. H105 is at the same position as H118 in L2a/b/c, while SME-1 has two residues different within the selected binding site (Figure 4C). The local environment of SME-1 is more hydrophobic due to the presence of Y104 at the loci of S117 in L2a/b/c and L167 at the loci of P180 in L2a/b/c. This difference is not indicated clearly in the model outputs, and we posit that this is due to the presence of the H105 in SME-1. This residue is closer to the centre of the active site, so it might influence a greater impact on the local features.

Meanwhile, KPC-2 altogether presents a different type of binding site environment, namely a more hydrophobic residue W105 is present at the loci of H118 of L2a/b/c and three other slightly different residues around the binding pocket.

The BindSiteS-CNN based binding site comparison results (represented as UMAPs) are consistent with the CVAE analysis, used both hydrophobic nodes and binding site residues as features. This suggests a correlation between local features of the binding site and global features of the six systems.

## Discussion

The over use of antibiotics in clinical practise has contributed to the emergence of β-lactamases that confer resistance. In spite of significant advancements in the molecular epidemiology and biochemical characterization of β-lactamases, the knowledge of the evolutionary forces driving the diversification of these enzymes remains limited. While most studies combine genetic, biochemical and structural analyses to assess the evolutionary processes that drive how β-lactamases confer resistance, they fail to specifically address the dynamics. This research presented here offers a comprehensive analysis of the L2 β-lactamase family as a representative case study.

The integrated sequence and structural alignment results, in conjunction with phylogenetic analysis, provide comprehensive insights into the evolutionary relationship and structural homogeneity of L2a, L2b, L2c, and L2d β-lactamase together with SME-1 and KPC-2. In the study of dynamics, the majority of findings indicate that computational models incorporating hydrophobic nodes and binding site residues effectively capture the varied dynamic behaviours of the L2 family, along with SME-1 and KPC-2, over long-time scales. The use of hydrophobic nodes in depicting dynamics is well-documented in protein folding, tertiary structure stability, and allostery (Galdadas *et al*, 2018, 2021; Olehnovics *et al*, 2021). This work introduces self-defined binding site residues to enhance the description of dynamics. The residues, characterized by relative conservation in sequence and structure, are crucial for maintaining the fundamental catalytic function of the class A β-lactamase core. Moreover, the results substantiate the reliability of these residues in representing the dynamics of class A β-lactamases. The hydrophobic nodes and binding site residues are pivotal in sustaining the conserved active site within the enzyme core of L2 β-lactamase variants. Such conservation is believed to contribute to the significant co-evolutionary information evident in both sequence and dynamic aspects. This underscores the importance of the kinetic properties of these residues in the evolutionary trajectory of β-lactamases. Environmental pressures may induce amino acid substitutions, potentially altering the conformation of aromatic residues. The conformational shifts are likely retained to modulate enzyme functionality (Liao *et al*, 2018). Observations indicate that both hydrophobic nodes and binding site residues aid in identifying these characteristics. Analyses of sequence, structure, and dynamics collectively reveal a consistent co-evolutionary pattern: KPC-2 → SME-1 → L2d → L2a/b/c This study employed hydrophobic nodes and binding site residues for CVAE modelling, capturing both global and local dynamics in the evolution of six β-lactamase systems. The dynamics were evident in the evolutionary trajectory from KPC-2 to SME-1 to L2d to L2a/b/c, highlighting the role of both global and local structural influences in the formation of active sites and the functional evolution of β- lactamases. Globally, a shift was observed where the majority of the Ω loop gradually distanced from the active site, while the α8 helix and adjacent residues approached the enzyme core. Concurrently, the core of the systems maintained stability due to the hydrophobic networks. Locally, active site residues increasingly converged, strengthening interactions, particularly with a key aromatic residue in the α3- α4 loop, which regulated access to the active site (Papp-Wallace *et al*, 2010; Olehnovics *et al*, 2021). These dynamic changes facilitated a functional shift in β-lactamase from carbapenemase to Extended-Spectrum Beta-Lactamase (ESBL) cephalosporinase activity, underscoring the Ω loop’s evolutionary significance. The BindingSiteS-CNN model, leveraging binding site information, emphasized detailed local interactions. The synergy of the poweful deep learning methods like CVAE and BindingSiteS- CNN, offer a comprehensive description of dynamics from local to global scales.

CVAE, with its unique architecture, adeptly handles complex dynamic 3-dimensional protein structures (Bhowmik *et al*, 2018). It enables precise clustering of protein conformations and associates each cluster with specific functional states, enhancing understanding of protein functions, dynamics, and biological phenomena (Akere *et al*, 2020; Olehnovics *et al*, 2021; Galdadas *et al*, 2018; Zhao *et al*, 2023; Cho *et al*, 2021). Meanwhile, BindingSiteS-CNN, utilizing a spherical representation-based graphical convolutional network, overcomes limitations of traditional 3D convolutional networks (Scott et al., 2022). This model excels in local protein environment similarity assessment, binding site classification, and predicting protein-ligand interactions, offering insights into physicochemical properties and biological functions across protein families.

Finally, the work presented here not only demonstrates the effectiveness of combining CVAE and BindingSiteS-CNN with MD in elucidating β-lactamase dynamics but also shows potential applicability to other protein families. It lays a foundation for deeper exploration into sequence-structure-dynamics-function relationships in class A β-lactamases. Furthermore, this integrated approach, combined with sequence and structure analysis, promises advancements in understanding enzyme-directed evolution.

## Materials and methods

### Structural Models

The crystal structures of L2b (PDB ID 5NE2) (Calvopiña *et al*, 2017), KPC-2 (PDB ID 3DW0) (Petrella *et al*, 2008), SME-1 (PDB ID 1DY6) (Sougakoff *et al*, 2002) were downloaded from the Protein Data Bank. The structures of L2a (UNIPROT ID P96465), L2c (UNIPROT ID P96465) and L2d (UNIPROT ID P96465) were obtained from the AlphaFold protein structure database (Jumper *et al*, 2021; Varadi *et al*, 2022).

### Sequence Alignment and Phylogenetic Tree Generation

Amino acid sequences of L2a, L2b, L2c, L2d, SME-1, and KPC-2 were derived from the structural files through systematic parsing of the respective PDB structure files. A multiple sequence alignment was conducted utilizing Clustal Omega version 1.2.4 (Madeira *et al*, 2022) employing its default parameters. The resulting alignment was subsequently visualized and interpreted with Jalview version 2.11.2.7 (Waterhouse *et al*, 2009) to furnish a graphical depiction of sequence congruities for improved comprehension. Building upon this, a phylogenetic tree was generated using iTOL version 6.8 (Letunic & Bork, 2021).

Given the scarcity of existing literature pertaining to the detailed L2 β-lactamase residues, a thorough annotation was conducted using UniProt BLAST (The UniProt, 2023) to determine the exact amino acid numbering for the four L2 systems. Consequently, the L2 β-lactamase sequence numbering was denoted using the structure of L2b β-lactamase (PDB ID 5NE2) (Calvopiña *et al*, 2017) while the SME- 1 and KPC-2 sequences were associated with PDB IDs: 1DY6 (Sougakoff *et al*, 2002) and 3DW0 (Petrella *et al*, 2008), respectively. Subsequent to this identification, an extended alignment of all six structures was conducted. Drawing upon the sequence and structural alignment of the hydrophobic nodes, coupled with the binding site residues delineated in KPC-2, the hydrophobic nodes and binding site residues in the remaining five systems were defined (Supplementary Table S1). Throughout this process, default parameters were consistently employed to ensure methodological reliability and the ensuing results informed next phases of data analysis and interpretation.

### Systems preparation and MD simulations

The initial system preparation utilized the PlayMolecule ProteinPrepare Web Application (Martinez-Rosell *et al*, 2017), where structural model files for the six systems were uploaded. Setting the pH at 7.4, heteroatoms were removed from the PDB files. ProteinPrepare autonomously executed pKa calculations and optimized hydrogen bonds, while simultaneously assigning charges and protonating the structure file within the high-throughput molecular dynamics (HTMD) framework (Doerr *et al*, 2016). The input files containing detailed information about atoms, bonds, angles, dihedrals, and initial atom positions were generated using tleap (Case *et al*, 2023) employing the Amberff14SB force field (Maier *et al*, 2015). Each system was solvated in TIP3P water model (Mark & Nilsson, 2001) within a cubic box, maintaining a minimum 10 Å distance from the nearest solute atom, and neutralized with 0.15 M Na^+^ and Cl^-^ ions. Prepared systems were initially minimized through 3000 iterations of steepest descent and subsequently equilibrated for 5 ns under NPT conditions at 1 atm. The temperature was steadily increased to 300 K with a time step of 4 fs, using rigid bonds, a 9 Å cutoff, and particle mesh Ewald summations (Cerutti *et al*, 2009; Wells & Chaffee, 2015) for long-range electrostatics. During equilibration, the protein backbone atoms were restrained, while the Berendsen barostat (Feenstra *et al*, 1999) controlled pressure and velocities were based on the Boltzmann distribution. The production phase consisted of multiple short MSM-based adaptively sampled simulations were run using the ACEMD engine (Doerr *et al*, 2016; Harvey *et al*, 2009). Each simulation was conducted in the NVT ensemble, used a Langevin thermostat with 0.1 ps damping and a hydrogen mass repartitioning scheme, allowing a 4 fs time step and recording trajectory frames every 0.1 ns. The MSM-based adaptive sampling algorithm utilizes multiple short parallel simulations and generates a discretised conformational state, which is then used to respawn further simulations. In this case, the MetricSelfDistance function was used to build the MSMs. In each round, 4 simulations of 50 ns each were run in parallel. The simulations were run until a minimum of 400 trajectories were obtained with each trajectory counting 500 frames and sampling a cumulative 20 μs for each system.

### Deep conformational clustering using CVAE

The utilisation of CVAE (Convolutional Variational AutoEncoders) was implemented in a systematic manner to investigate the evolutionary dynamics of six β-lactamase systems, namely L2a, L2b, L2c, L2d, SME-1, and KPC-2. Data derived from each protein system, including the root-mean-square deviation (RMSD), contact, and radius of gyration, were computed using MDAnalysis (Michaud-Agrawal *et al*, 2011) and MDTraj (McGibbon *et al*, 2015). Detailed analyses involved the formulation of pairwise distance maps extracted from every fifth frame of the 250 trajectories in each system. The focus was directed towards hydrophobic nodes and binding site residues featuring the constraint of Cα atom distances ≤8 Å, which were recorded as non-zero values in a specified three-dimensional matrix.

The cumulative data, represented as 64 × 64 distance matrices, were consolidated into a unified 3D matrix for every system, accompanied by a label file containing pertinent metadata. The python code for model implemented in the current study was adopted from Bhowmik et al. (Bhowmik *et al*, 2018) The CVAE’s encoding section was structured with an 80:20 validation ratio and underwent training across 100 epochs. Dimensions spanning from 3 to 30 were explored, eventually settling on the 21st dimension, which exhibited the minimal loss, for the model’s architecture. Continuous oversight was maintained for potential overfitting. For the decoding component, matrices and label files derived from four L2 systems were used to assess the model’s performance and to discern the clustering patterns of the conformations inherent to these systems. Additionally, data from SME-1 and KPC-2 were incorporated into another distance matrix and labels to observe dynamic shifts occurring through the evolutionary pathway of β-lactamase.

The UMAP algorithm was then employed to reduce the dimensionality of the decoded embedding into two dimensions, thereby simplifying visualization. This, when combined with the free energy landscape, proved instrumental in isolating distinctive conformations from energetically favourable regions. The CVAE workflow can be fetched in Figure S5 and the script operationalized for CVAE was a modified version of a pre-existing in-house code.

### BindSiteS-CNN based binding site comparison

BindSiteS-CNN was employed to capture the differences between the active site local features of the six systems: L2a, L2b, L2c, L2d, KPC-2 and SME-1. The methodology encompassed binding pocket surface preparation and BindSiteS-CNN model processing (Figure S6) and was adopted from Scott et al. (Scott *et al*, 2022). The samples were taken every fifth frame from those used for CVAE.

In the binding pocket surface preparation phase, the binding pocket surface of each frame was generated with side chain atoms of the binding site residues as the filtering reference. The computed pocket surface meshes with vertices enriched with physicochemical information describing the hydrophobicity, electrostatic potential and interaction-based classification of surface-exposed atoms lining the pocket were saved as PLY files and integrated as part of an in-house β-lactamases active pocket database.

During the BindSiteS-CNN model processing stage, the prepared 3D pocket mesh objects were fed into the trained BindSiteS-CNN model as input data. UMAP has been used to visualize their distribution in the descriptor space, those with similar binding sites would cluster together.

### Structural Analysis

The trajectories of the molecular simulations were meticulously aligned to their corresponding reference structures with MDAnalysis (Michaud-Agrawal *et al*, 2011) and MDTraj (McGibbon *et al*, 2015). The stride of frames within these trajectories was retrieved using the identical set of tools. To elucidate the general dynamics features inherent in the trajectories, calculations were performed again leveraging the functions within the MDTraj and MDAnalysis packages. For a more visual and intuitive understanding, the trajectories were loaded into the Visual Molecular Dynamics (VMD) software (Humphrey *et al*, 1996)h. This tool also facilitated the superimposing of structures and enabled a comprehensive conformational comparison. After delineating the spatial variations between distinct conformations, visual representations were generated via the Protein Imager (Tomasello *et al*, 2020). Additionally, the Matplotlib package (Hunter, 2007) in Python was employed for all statistical and graphical representations, including plots and figures, to present the data in a comprehensive and interpretable manner.

### Data Availability

All input files to run the simulations and the molecular dynamics trajectories can be downloaded from DOI 10.5281/zenodo.10500538

## Author Contributions

**Yu Zhu:** Formal analysis, validation, investigation, visualisation, Writing - original draft; **Jing Gu:** Formal analysis, validation, investigation, visualisation, writing original draft; **Zhuoran Zhao:** Running adaptive bandit simulations; Data curation **Edith A W Chan:** Methodology, Software; **Maria F Mojica:** Conceptualization, Writing - review and editing **Andrea M Hujer:** Writing – reviewing and editing; **Robert A Bonomo:** Conceptualization, Supervision, Writing – review and editing; **Shozeb Haider:** Conceptualization, Methodology, Supervision, Validation, Writing - review and editing,

## Acknowledgements

This work was supported in part by funds and/or facilities provided by the Cleveland Department of Veterans Affairs, Award Number 2I01BX001974 (R.A.B.), from the Biomedical Laboratory Research & Development Service of the VA Office of Research and Development. The content is solely the authors’ responsibility and does not necessarily represent the official views of the Department of Veterans Affairs.

