## Supplementary Information for "Deciphering the co-evolutionary dynamics of L2 β-lactamases via Deep learning"

**Table S1. Feature selections for the six class A β-lactamase.**

| System | Numbering Reference | Hydrophobic Nodes | Binding Site Residues |
|---|---|---|---|
| **KPC-2** | PDB ID: 3DW0 | G74, F75, L76, A77, A78, A79, V80, L81, A82, A101, L102, V103, S106, P107, I108, A124, A125, A126, V127, A133, A134, A135, L137, L138, L139, A185, V186, T187, A198, L199, A200, A201, A223, A224, V225, A230, V231, G232, A248, V249, V250, V260, L261, A262, A280, A281, A282, A283 | S70, K73, P104, W105, P107, S130, N132, E166, L167, N170, T216, K234, T235, G236, T237, C238, G239 |
| **SME-1** | PDB ID: 1DY6 | G74, F75, L76, A77, A78, A79, V80, L81, E82, D101, L102, E103, S106, P107, I108, S124, A125, A126, L127, G133, A134, T135, I137, I138, M139, A185, V186, A187, V198, L199, N200, A201, A223, S224, V225, V230, V231, G232, A248, V249, I250, I260, V261, S262, A280, E281, A282, S283 | S70, K73, Y104, H105, P107, S130, N132, E166, L167, N170, T216, K234, T235, G236, S237, C238, G239 |
| **L2a** | UniProt ID: P96465 | S87, V88, L89, A90, A91, T92, V93, L94, S95, D114, L115, L116, A119, P120, V121, R137, A138, T139, I140, T146, A147, A148, L150, L151, F152, A198, M199, A200, V211, L212, Q213, L214, A236, G237, L238, R243, V244, R245, A260, V261, L262, V282, L283, T284, A292, Q293, V294, G295 | S83, K86, S117, H118, P120, S143, N145, E179, P180, N183, T229, K247, T248, G249, S250, N251, G252 |
| **L2b** | PDB ID: 5NE2 | S87, M88, L89, A90, A91, T92, V93, L94, S95, D114, L115, L116, A119, P120, V121, R137, A138, T139, I140, T146, A147, A148, L150, L151, F152, A198, M199, A200, V211, L212, Q213, P214, A236, G237, L238, R243, V244, G245, A260, V261, L262, V282, L283, T284, A292, Q293, V294, G295 | S83, K86, S117, H118, P120, S143, N145, E179, P180, N183, T229, K247, T248, G249, S250, N251, G252 |
| **L2c** | UniProt ID: P96465 | S87, V88, L89, A90, A91, T92, V93, L94, S95, D114, L115, L116, A119, P120, V121, R137, A138, T139, I140, T146, A147, A148, L150, L151, F152, A198, M199, A200, V211, L212, Q213, P214, A236, G237, L238, R243, V244, G245, A260, V261, L262, V282, L283, T284, A292, Q293, V294, G295 | S83, K86, S117, H118, P120, S143, N145, E179, P180, N183, T229, K247, T248, G249, S250, N251, G252 |
| **L2d** | UniProt ID: P96465 | S88, M89, L90, V91, A92, H93, V94, L95, S96, D115, L116, L117, A120, P121, V122, R138, G139, T140, L141, T147, A148, A149, L151, L152, L153, A199, M200, A201, G212, L213, Q214, F215, A237, G238, L239, R244, V245, G246, A261, V262, L263, V283, V284, T285, A293, R294, V295, G296 | S84, K87, S118, Y119, P121, S144, N146, E180, P181, N184, T230, K248, T249, G250, S251, N252, G253 |

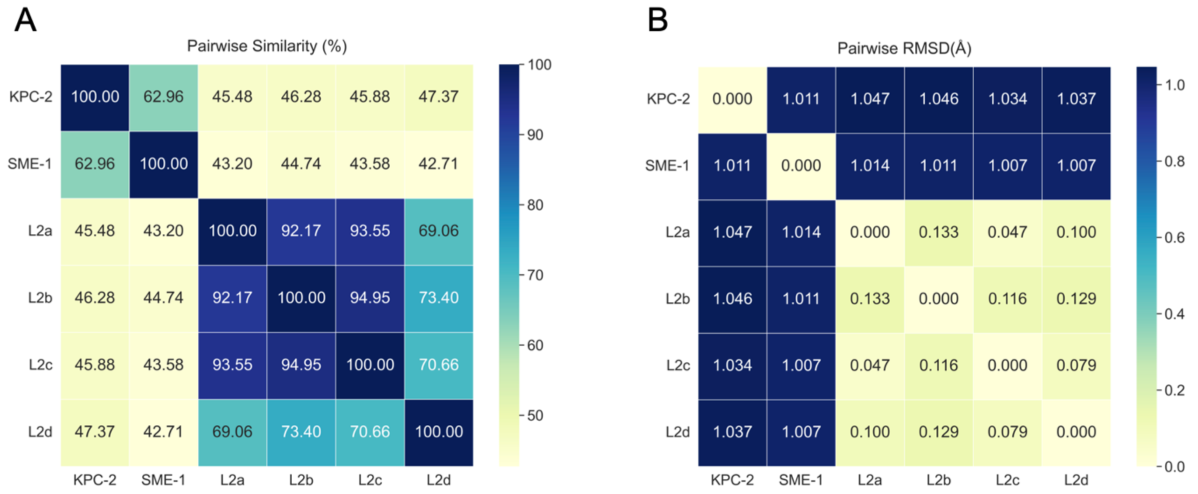


**Figure S1. Sequence and structure alignments.** (A) The pairwise sequence similarity of the six class A β-lactamases. (B) The pairwise structural Cα RMSD of the six systems.

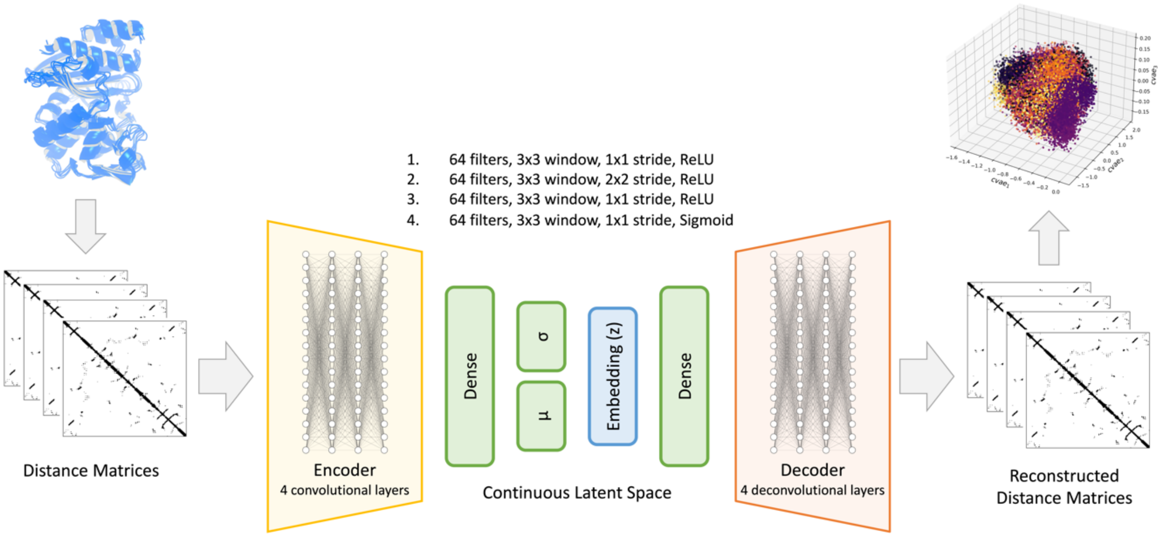


**Figure S2. Core architecture of the CVAE-based deep learning approach.** Convolutional Variational Autoencoder (CVAE) is a type of artificial neural network that employs the principles of both Convolutional Neural Networks (CNNs) and Variational Autoencoders (VAEs). It uses convolutional networks as its encoder and decoder and leverages the principles of variational inference to learn a structured representation of the input, enabling the generation of new, similar data. Simulation trajectories and feature selections are required for the model training. The decoded embeddings can be processed via diverse dimension reducing methods. The CVAE provides a holistic view of protein structures and their dynamic changes, enabling more precise clustering of different protein conformations. This ability to capture and cluster subtly different protein conformations has applications across various areas, from advancing our understanding of diseases linked to protein misfolding to facilitating drug discovery by mapping potential drug-binding pockets in different conformational states.

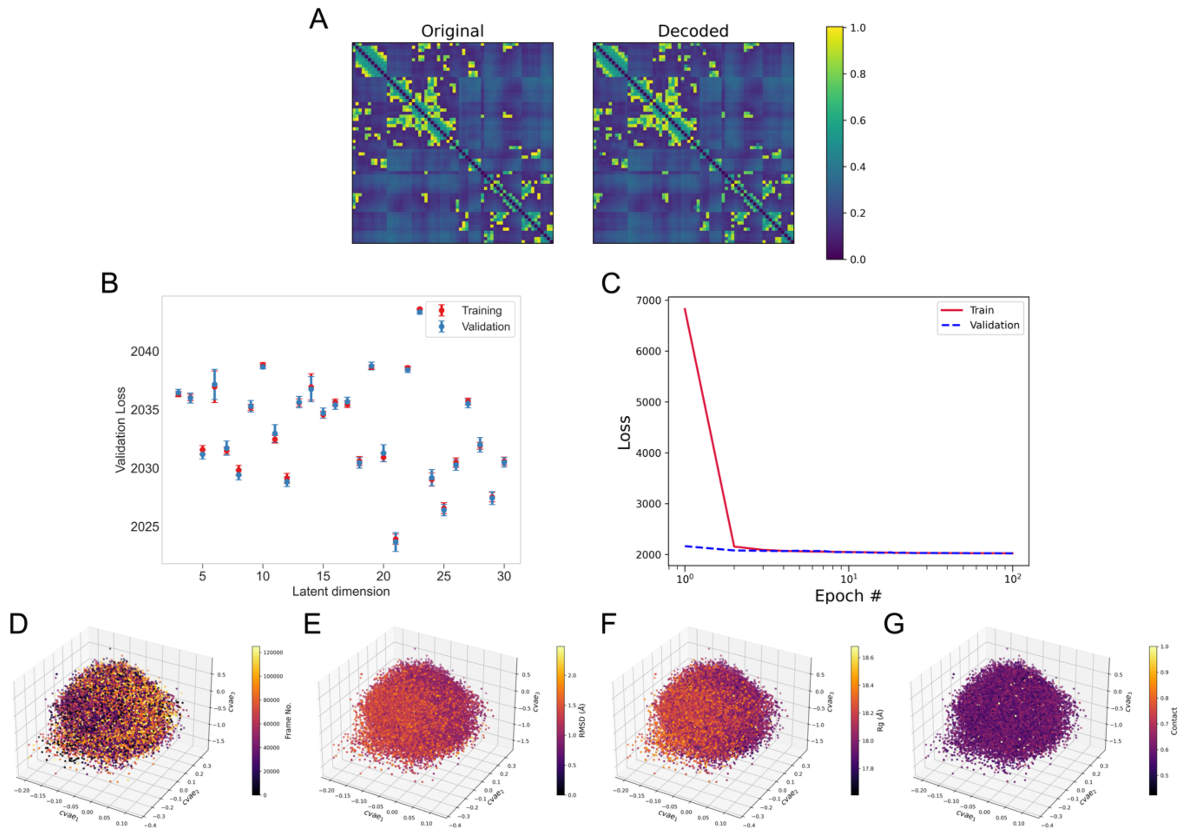


**Figure S3. CVAE performance using the four L2 β-lactamases trajectories.** (A) Comparison between the original distance matrix and the decoded matrix. (B) Loss for each latent dimension. (C) Model training and validation for the 21st latent space dimension for each epoch. (D), (E), (F) and (G) represent the 3D embeddings of the stacked systems in frame number, RMSD, radius of gyration, contact, respectively.

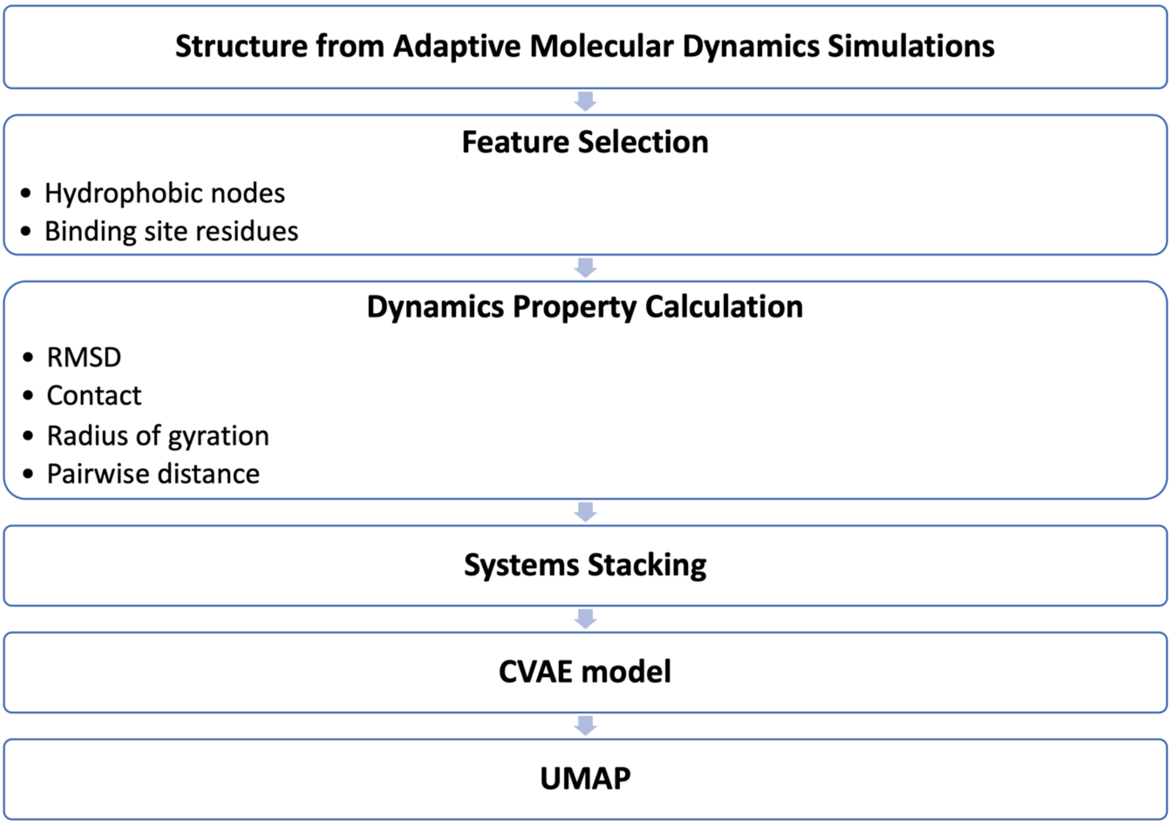


**Figure S4.** Flowchart of CVAE based clustering.

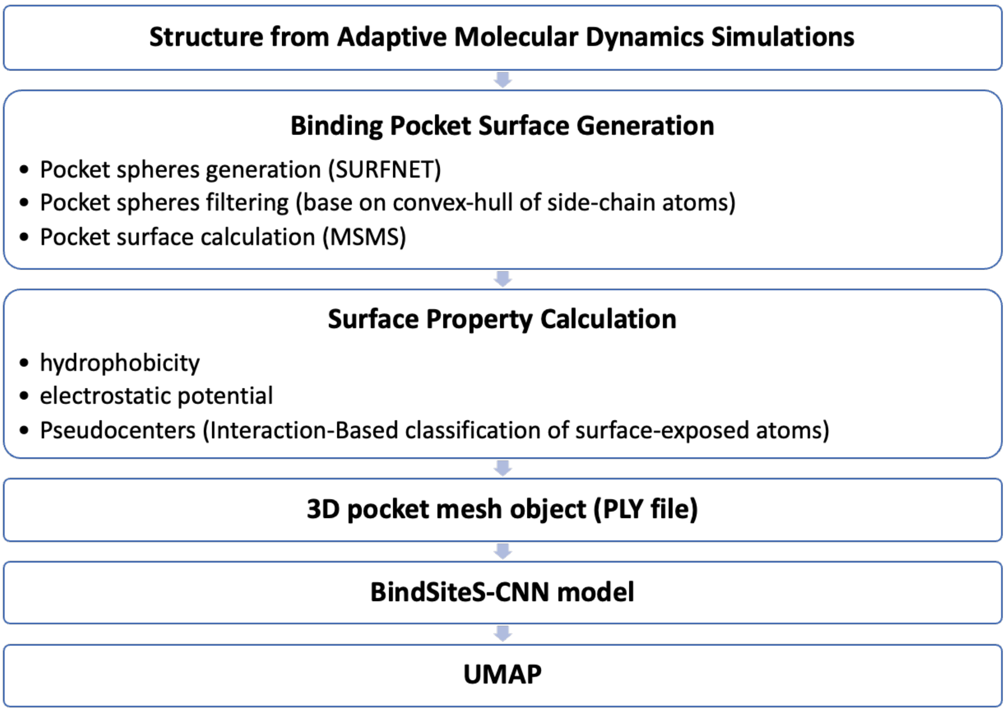


**Figure S5.** Flowchart of BindSiteS-CNN based binding site comparison.
